# Rock the Chalk: A five-year comparative analysis of a large microbiology lecture course reveals improved outcomes of chalk-talk compared to PowerPoint

**DOI:** 10.1101/644567

**Authors:** Christopher M. Waters

## Abstract

The rise of electronic assisted presentation programs such as PowerPoint in undergraduate large lecture biology classes has displaced more traditional hand-drawn lectures such as the blackboard or overhead projectors, referred here as “chalk-talk” approaches. But which method is more effective in a large lecture microbiology classroom is unclear. Here I present data from a large microbial genetics lecture course taken during a five-year span comparing PowerPoint to chalk-talk lecturing methods. The results indicate that the chalk-talk approach was preferred by the students and rated higher in all measured metrics including course enjoyment, learning of key concepts, and course outcomes.

## Introduction

Since the introduction of electronic assisted learning, presentation programs such as PowerPoint have become the mainstay of undergraduate biology education. However, it is not clear whether this method of teaching is superior to past methods such as the use of blackboard or overheads in which the instructor writes/draws lectures on a blank canvas, approaches that I will refer to as “chalk-talk”. The studies addressing this question have arrived at different conclusions regarding the effectiveness of each presentation style, making it unclear which method is more effective. For example, Susskind determined that using PowerPoint in an Introduction Psychology course did not impact performance but did increase student attitudes (1). This study, which utilized two different classes, delivered half of the lectures by traditional chalk-talk while the other half were delivered using PowerPoint, with the lectures counterbalanced in each class to control for differences in the topics presented. Alternatively, Savoy et. al. reported that students maintained 15% more of the verbal information given by the instructor using the chalk-talk approach for a given lecture, even though the students indicated a preference for PowerPoint (2). A similar study in a medical physiology course that examined one lecture presented in both styles supported Susskind’s conclusion as no significant difference in outcomes between chalk-talk versus PowerPoint was observed, although the students reported significant differences in learning metrics between the two approaches. For example, in post-class surveys the students indicated that PowerPoint was better for “clarity of words” and “summarizations” while chalk-talk was superior for “clarity of concepts”, “learning to draw diagrams”, and “understanding the subject” (3). Similar results were obtained by Kumar comparing PowerPoint to chalk-talk for 5^th^ year surgery students using post-class surveys (4). PowerPoint ranked higher for content and clarity while chalk-talk ranked higher for stimulating interest and advancing understanding, although the difference for this last category was minimal (4). Bartsch et. al. found that in a social psychology course PowerPoint slides with both images and text decreased performance compared to overheads (5). This same report concluded that inclusion of non-related images on a PowerPoint slide was detrimental in retaining the concept presented. The bottom line is that multiple studies exist that show a student preference for PowerPoint over chalk talk (6,7) or vice versa (8,9), and there is not clarity to which method is a more effective teaching tool.

Why is there such a discrepancy in studies that compare PowerPoint to chalk-talk? I postulate although the studies cited above were well-controlled, it is difficult to compare the two methodologies in this manner because these lectures/courses were not designed around a given presentation format. More importantly, there is a learning curve for the instructor to effectively deliver a lecture using PowerPoint or chalk-talk and comparing these methods with only one or a subset of lectures does not allow for instructor mastery of the different approaches. Therefore, instructor skill in each approach could lead to different outcomes. Thus, the results of studies comparing these approaches are mixed (10).

I am aware of only one study addressing the effectiveness of PowerPoint versus chalk-talk in my own field of microbiology, and it concluded that although students preferred PowerPoint, use of a blackboard was the most effective; however, the methodology for assessing effectiveness is not clear (11). Here, I harness my own experiences teaching the large lecture course “Microbial Genetics” at Michigan State University for the last ten years, the first seven of which utilized PowerPoint and the last three that utilized chalk-talk, to analyze the effectiveness of PowerPoint versus chalk-talk in microbiology education at the undergraduate level. An analysis of course metrics including course reviews, course attendance, and student assessment indicates that the chalk-talk approach was superior to PowerPoint in every category.

## Methods

### Course structure

The data are gathered from five semesters of “MMG 431:Microbial Genetics” given in the fall semester at Michigan State from 2013 to 2017 in which I was the sole instructor. PowerPoint was exclusively used as the lecture format in 2013 and 2014 while a chalk-talk approach was primarily used in 2014-2017. Course enrollment for these years in parentheses is 2013 (123), 2014 (151), 2015 (155), 2016 (134), and 2017 (150), and it was taught in a lecture hall that seats about 200 students. The course met three times a week for 50-minute lectures. Students were evaluated by three midterm exams consisting of 53 multiple choice questions and a cumulative final exam consisting of 95 multiple choice questions covering key concepts from throughout the semester.

### Lecture format

During 2013 and 2014, PowerPoint slides were projected onto two screens at the front of the class, and one lecture had on average 20-25 slides. An electronic file of the PowerPoint slides was provided to the class before the lecture. The class was asked ∼2-4 “think:pair:share” questions every lecture, which has been demonstrated to increase cognitive learning in large lecture classroom (12-14). For the chalk-talk approach, a touch screen laptop was utilized and projected to the same two screens, and lectures were hand-drawn with an electronic stylus using the program OneNote. Using OneNote rather than a traditional blackboard better projected the chalk-talk lectures, allowed the use of multiple colors, and enabled rapid switching to other programs such as PowerPoint, iClicker software, or a web browser. The lecture started as a blank white page, and text and pictures were drawn with a touch screen stylus using various line widths and colors. Each lecture was a new OneNote page that was a subset of an individual folder so all lecture notes were saved and could be referenced later. The lecture notes were not distributed to the class. PowerPoint was used sparingly (∼0-2 slides per class) to show complex diagrams, structures, or more commonly (∼2-4 slides per class) to project the same “think:pair:share” questions that were used for the PowerPoint lectures. As the intention during these five years was to provide the best learning experience and not execute a tightly controlled study, there were modifications in course structure and procedures from year to year. For example, on-line homework was assigned in 2014-2016, but not in 2013 and 2017. In 2017, the audio of all lectures was recorded and posted on the central course page allowing students to re-listen to lectures (but no lectures notes were provided).

### Student course evaluation

At the end of the semester, students are asked to complete an anonymous 21-question Student Instructional Rating System (SIRS) evaluation form that is standard for all classes at Michigan State University (Table 1). Students can opt out of completing these evaluations, and it is not known how many students completed them for each year. These questions are scored from 1 to 5 using the Likert scale of: (1) Superior (2) Above Average (3) Average (4) Below Average and (5) Inferior. The questions are then aggregated into the six general categories listed in Table 1, and a mean score for each category is determined. Students can also submit specific comments about the course although this is optional. During the five years analyzed here, the percentage of class that submitted a specific comment was 2013 (31.7%), 2014 (25.8%), 2015 (37.4%), 2016 (23.9%), and 2017 (21.3%). These comments were binned into three categories of positive (mostly positive comments), neutral (an equal mix of positive and negative comments), or negative (mostly negative comments) by four individuals that are not connected to this study or the course without the evaluators knowing the year the comment was made.

**Table 1:** Questions on student Instructional Ratings Systems Evaluation Forms.

| **Category** | **Question** |
| --- | --- |
| Instructor Involvement | The instructor's enthusiasm when presenting course material. |
| Instructor Involvement | The instructor's interest in teaching. |
| Instructor Involvement | The instructor's use of examples or personal experiences to help get points across in class. |
| Instructor Involvement | The instructor's concern with whether the students learned the material. |
| Student Interest | Your interest in learning the course materials. |
| Student Interest | Your general attentiveness in class. |
| Student Interest | The course as an intellectual challenge. |
| Student Interest | Improvement in your competence in this area due to this course. |
| Student Instructor Interaction | The instructor's encouragement to students to express opinions. |
| Student Instructor Interaction | The instructor's receptiveness to new ideas and others' viewpoints. |
| Student Instructor Interaction | The student's opportunity to ask questions. |
| Student Instructor Interaction | The instructor's stimulation of class discussion. |
| Course Demands | The appropriateness of the amount of material the instructor attempted to cover. |
| Course Demands | The appropriateness of the pace at which the instructor attempted to cover the material. |
| Course Demands | The contribution of homework assignments to your understanding of the course materials relative to the amount of time required. |
| Course Demands | The appropriateness of the difficulty of assigned reading topics. |
| Course Organization | The instructor's ability to relate the course concepts in a systematic manner. |
| Course Organization | The course organization. |
| Course Organization | The ease of taking notes on the instructor's presentation. |
| Course Organization | The adequacy of the outlined direction of the course. |
| Enjoyment | Your general enjoyment of the course. |

### Course participation

During all five years, students were given 2-4 active learning “think:pair:share” exercises during the lecture. These exercises consisted of a question proposed to the class, followed by the class dividing into groups for ∼1 minute to discuss. Each student then used an iClicker to submit a response, and the question and correct answered were discussed as a class. Students were assigned participation points based on how many responses they submit, and credit was not dependent on selecting the correct answer. iClicker responses were used to measure student attendance as presented in Figure 4.

### Ethical considerations

This study was reviewed by the Michigan State University Institutional Review Board and found to be exempt under 45 CFR 46.101(b) 4. All data used is anonymous and cannot be linked to individual students.

## Results

### Rationale for implementing chalk-talk lectures

I have been the course administrator and sole instructor for the junior/senior level undergraduate course “MMG 431:Microbial Genetics” at Michigan State University since 2013, and I co-taught this course with one other instructor from 2009-2012. Therefore, for this study I only analyze data from 2013-2017 where I was the only instructor and course administrator. This course is taught once a year with student enrollments ranging from 121 to 155, and it is required for graduation with a degree in Microbiology. From 2009-2014, I had utilized a standard lecture course format with PowerPoint slides interspaced with active learning “think:pair:share” exercises. The students were given access to printable PDFs of all lecture slides before class, and these were intended to be an outline for further note taking during class. Because the lecture slides were available, many students preferred to bring their laptop computers and followed the lecture electronically. From my perspective, many of the students were not attentive during the lecture. Visitors to the course also noted that students often participated in non-class related activities on their electronic devices such as social media. Students expressed both to me in person and on in-class evaluations that too much material was covered too rapidly. For example, this comment from 2014 is representative of many student comments during the years when PowerPoint was used: “I feel that the appropriateness of the amount of material the instructor attempted to cover and the pace at which the instructor attempted to cover the material was too great.”

To attempt to address both of these problems, in 2015 I stopped primarily lecturing with PowerPoint and rather presented the lecture in a chalk-talk style. Using this approach, I wrote the information as I lectured, and the students were required to take notes in real time. The students were not provided any written notes either before or after class, and thus they were required to actively take notes and pay attention during the lecture. I did continue to implement 2-4 think:pair:share exercises during class, and the students received participation points for each response regardless of whether they were correct. On occasion, complicated structures were shown as PowerPoint slides, although this was rarely greater than 1 slide per lecture and many lectures presented no information using PowerPoint. As this course was taught by the same instructor teaching the same general material, with two years of PowerPoint (2013-14) and three years of chalk-talk (2015-17), an analysis of course data provides an excellent opportunity to gauge which approach was more effective in a microbiology large lecture course.

### The instructor perspective

I found the students were clearly more engaged when lecturing using chalk-talk and few actually used laptops or other electronic devices during class. The students asked more questions during the lecture and more actively participated when discussing think:pair:share activities. Another advantage was that lecturing using the chalk-talk style allowed more flexibility than PowerPoint. For example, if a student asked a question about a topic that was not in the original lecture, I was able to switch directions mid-class without being constrained by the next slide. I would often think of new think:pair:share questions during the lecture and could stop and present these to the class, which was not feasible using only PowerPoint. Essentially, my perspective was that lecturing using chalk-talk is a much more active and engaging experience for both myself and the students than using PowerPoint, and I was better able to connect with the class.

### Student course reviews indicate students prefer chalk-talk over PowerPoint

To assess student perception of the two lecture formats, student responses to the 21 questions grouped into six categories, as listed in Table 1, from SIRS evaluation forms from 2013-2017 were assessed. The scores for these six categories from 2013 to 2014 where PowerPoint lecturing was used were quite consistent with scores ranging from 1.95 to 2.88 graded on the Likert scale with (1) Superior (2) Above Average (3) Average (4) Below Average and (5) Inferior (Fig. 1A). Starting in 2015, when chalk-talk was implemented, the scores of every category improved annually. In 2017, the scores ranged from 1.37-1.8. In fact, the lowest numeric value for each category was measured in 2017 (Fig. 1A). In 2013 and 2014, only the “Student/Instructor” category was less than 2, but all six categories were below 2 in 2017. One possibility to explain these results is that I simply improved generally as an instructor over time; however, this hypothesis is unlikely for several reasons. First, 2013 was my fifth year teaching this course so I was already highly experienced with the PowerPoint approach and the course material. Second, the scores from 2013 to 2014 are stable, and they are consistent with SIRS scores from 2009-2012. In fact, half of the categories had worse scores in 2014 than in 2013. Improvement was only observed in 2015 when the format was changed to the chalk-talk style, and this improved every year as I became more adept at lecturing using this method.

**Fig. 1.**
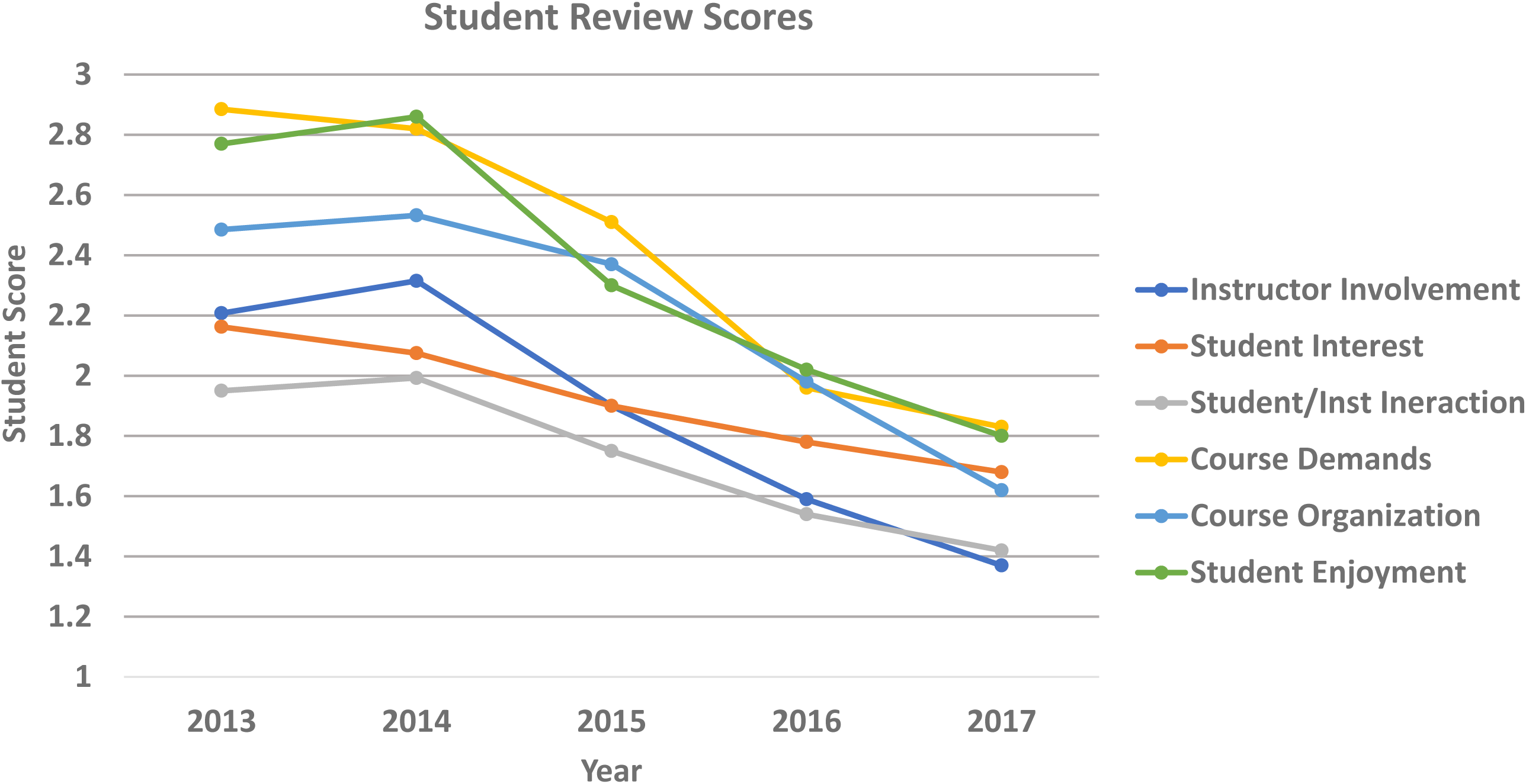
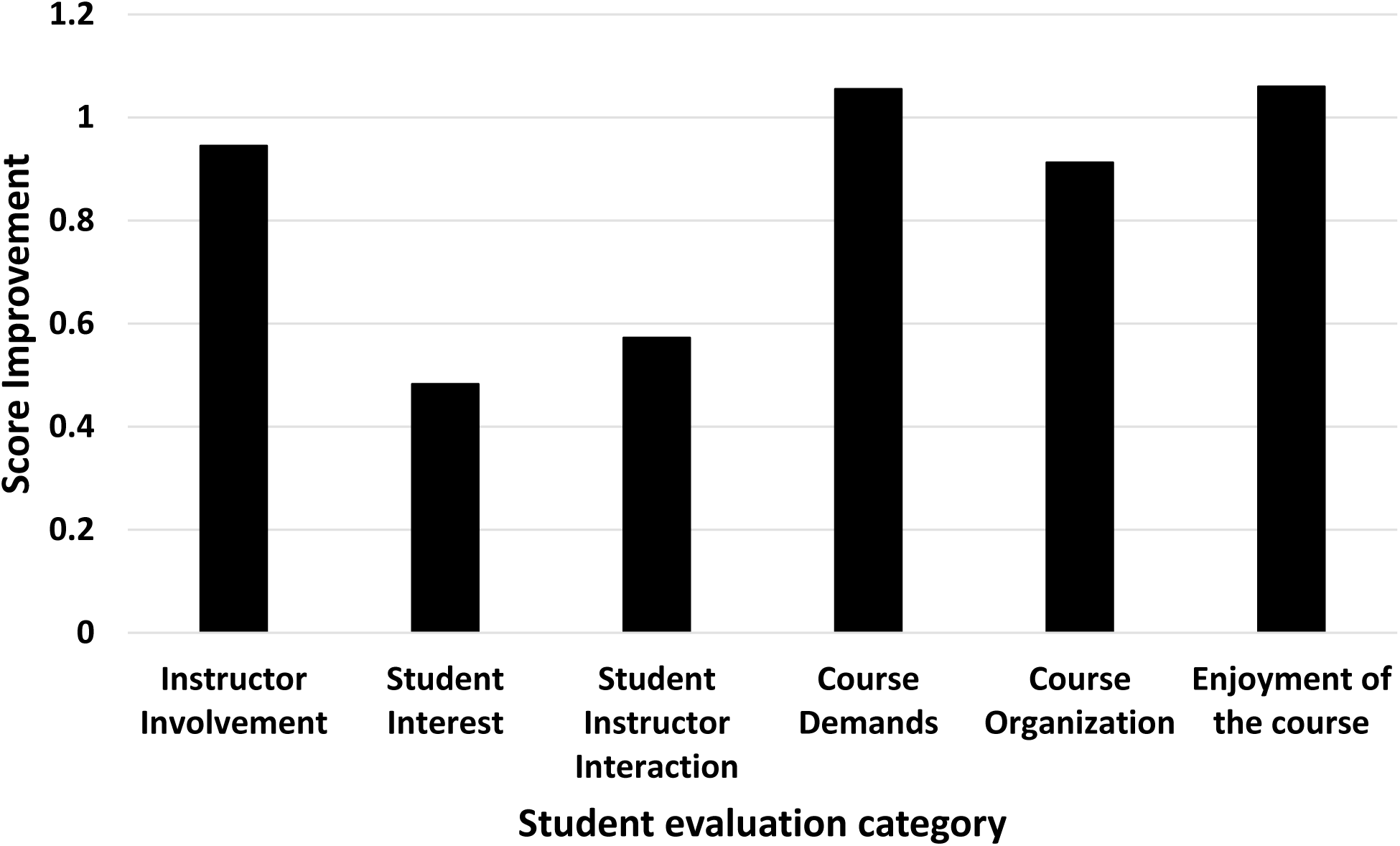
Analysis of student evaluations. Student evaluation responses from 2013-2017 to the six categories described in Table 1 were analyzed. (A) Student evaluations are grouped into six general categories, and the score from each year is shown with (1) Superior (2) Above Average (3) Average (4) Below Average and (5) Inferior. (B) The maximum increase in scores for each group during this five year period is shown.

To determine which category exhibited the greatest improvement, I subtracted the lowest scores (all in 2017) from the highest scores (either in 2013 or 2014) in each category (Fig. 1B). “Enjoyment of the Course” and “Course Demands” showed the greatest gain in student evaluation scores increasing by over 1 point followed by “Instructor Involvement” at 0.95, but all categories exhibited increases greater than 0.48.

The final data that was analyzed from the SIRS evaluations, which is perhaps the most informative, is taken from the specific comments section. Four individuals not connected to this study independently analyzed the specific comments provided by the students for all five years and binned them into positive (i.e. no negative comments), negative (i.e. no positive comments), or neutral categories (i.e. a mix of positive and negative comments). Specific comments were not required to complete the form and ranged from 29-56 entries per year. The mean percentage and standard deviation for each category of the four evaluators per year is shown in Fig. 2. In 2014, negative comments were significantly greater than either positive or neutral comments, but this switched when chalk-talk was implemented as positive comments were significantly greater than neutral or negative comments from 2015-2017. Neutral comments also declined during this time. Positive comments from 2015-17 expressed enthusiasm for the chalk-talk style with a representative comment being:

**Fig. 2.**
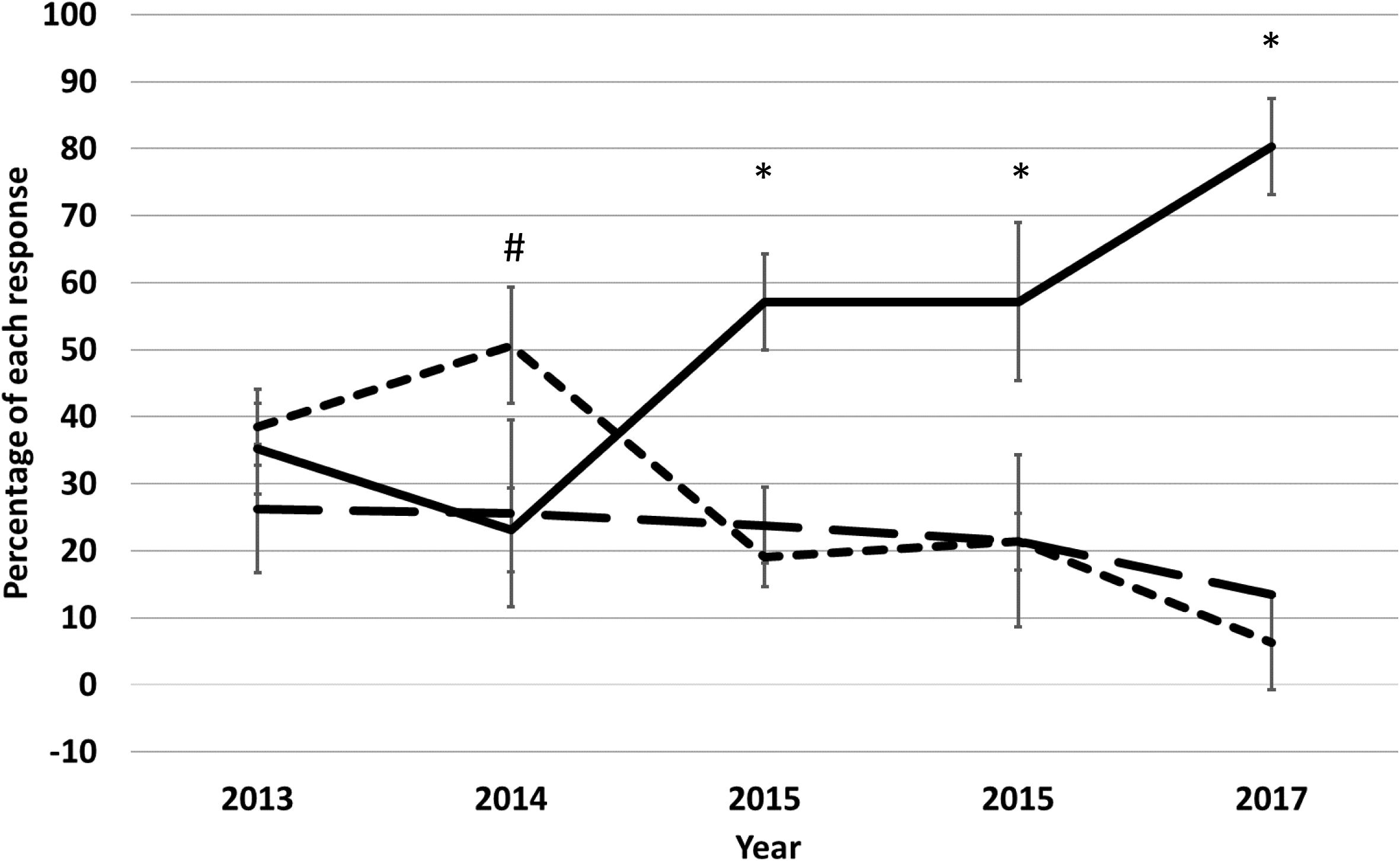
Student SIRS comments between PowerPoint and chalk-talk. The specific student comments from every student evaluation was grouped as either positive, neutral, or negative by four individuals and calculated as a percentage of the total comments. Solid=positive, large dashes=neutral, small dashes=negative. The mean percentage with the standard deviation is plotted for each year. #-Negative versus positive and neutral (p<0.0002), *-Positive versus negative and neutral (p<0.0001) as determined by a Two-Way ANOVA followed by Tukey’s Multiple Comparison Test.

> “His way of lecturing is so great. I wish it would be mandatory for every single class at MSU to mimic his teaching style. He writes down his notes and discusses them with us. it shows us if he can write it down we can too. It’s not too fast but we learn a lot.”

### Chalk-talk improves grades

To assess the difference in course outcomes between the two lecture formats, I analyzed course grades across each year. The percentage of the class that received the highest score of 4.0 versus a failing grade of 0 is plotted for each year (Fig. 3). As seen with the SIRS scores, these percentages were quite similar between 2013 and 2014 when PowerPoint was used, and these are reflective of the course from 2009-2012. However, students receiving the highest grade of a 4.0 showed a consistent increase from 2015 to 2017 reaching a total of 40% of the class in 2017. The large increase in 4.0 scores from 2016 to 2017 may be attributed to the posting of audio recordings of the lecture, nevertheless, 2015 and 2016 both have a higher percent of students receiving 4.0 than 2013 or 2014. Conversely, students receiving the lowest grade of 0 decreased in 2015 to 2017. Students who failed the course dropped from 17.2% in 2013 to 2.7% in 2017. A similar decrease was observed for lower scores of 1.0 and 1.5, but mid-level grades of 3.5, 3.0, 2.5, and 2.0 remained constant during this time frame (data not shown).

**Fig. 3.**
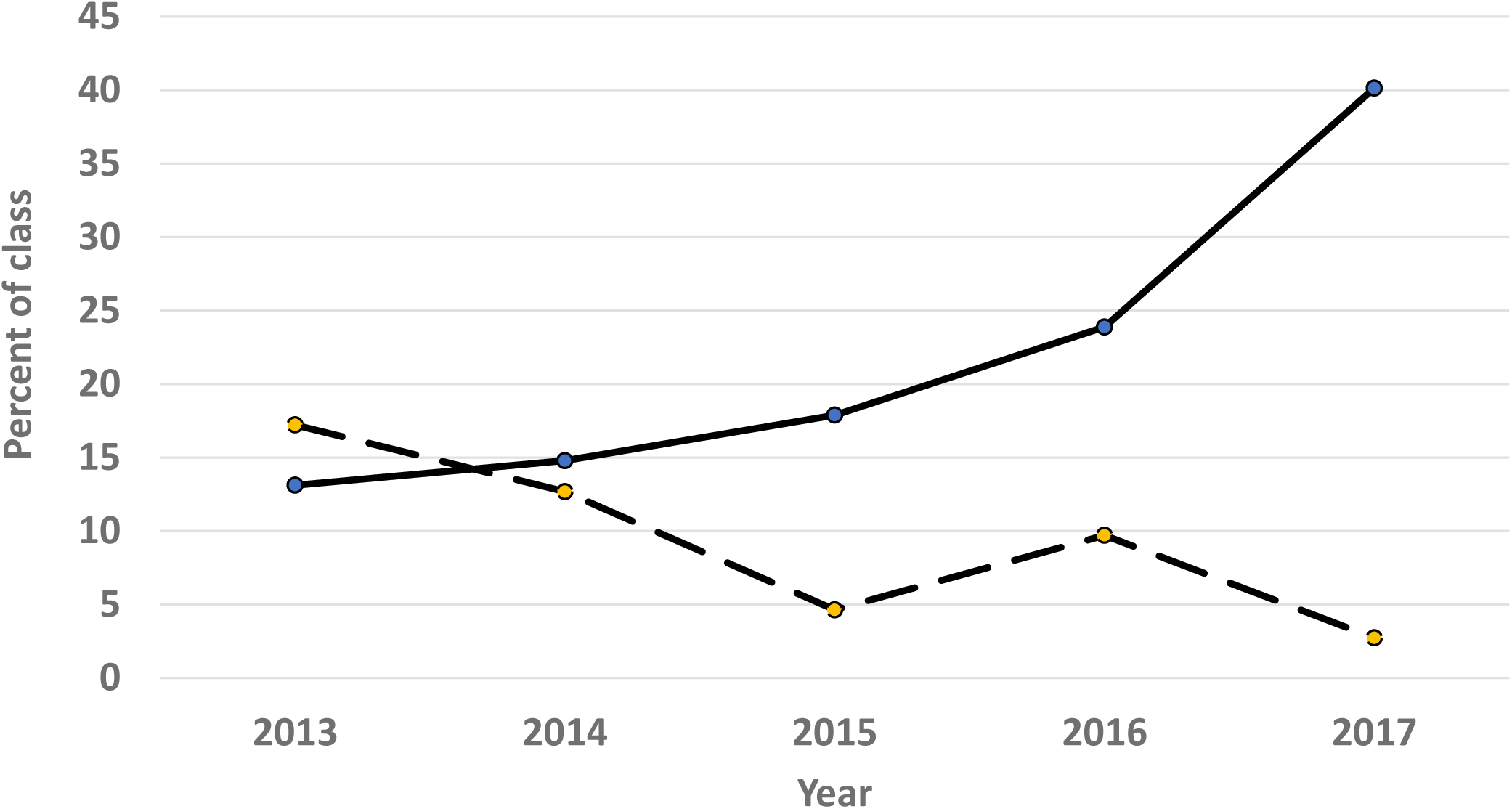
Student grades increase with chalk-talk methodology. Grades are assigned on a 4.0 scale, and the percent of the class that would receive a 4.0 (solid) versus a failing 0 (dashed) is shown each year.

Analyzing score adjustments for final grades also highlights the effectiveness of the chalk-talk approach. The data shown in Fig. 3 are the scores that would have be assigned based on the percentage of points received, but in 2013 and 2014 small but equal increases were given to all students to adjust for course difficulty. No such bumps were given in 2015-2017 as the raw scores were improved such that this grade adjustment was no longer necessary.

### Course attendance

Because students are no longer provided lecture notes, I hypothesized that course attendance would increase. Course attendance can be assessed by analyzing the average percentage of iClicker points obtained for the class for responses to in-class questions. These results show increased participation when chalk-talk was used as the three highest participation scores for these five years are from 2015-2017 (Fig. 4).

### Learning of key concepts

Switching to chalk-talk certainly reduced the amount of material that I could cover, as discussed below, which could account for the changes observed. However, it is important to note that my syllabus, including the key concepts and dates they would be covered, was unchanged from 2013-2017. To assess retention of key concepts at the completion of the course, I analyzed the results of the final exams from 2013-2017. These exams are cumulative, and they test the students on key concepts from the entire course. Because the final exam focuses on key concepts, and less on individual details, these exams were highly similar from 2013-2017 and can serve as a standard to compare concept retention. The highest median final exam scores occurred in 2015-2017 when chalk-talk was used while the lowest median final exam scores were 2013-2014 when PowerPoint was used (Fig. 5A). This increase in final exam scores occurred even though many students had less motivation to study for the final in 2015-2017 because they had obtained higher grades on the first three midterm tests throughout the semester.

**Fig. 4.**
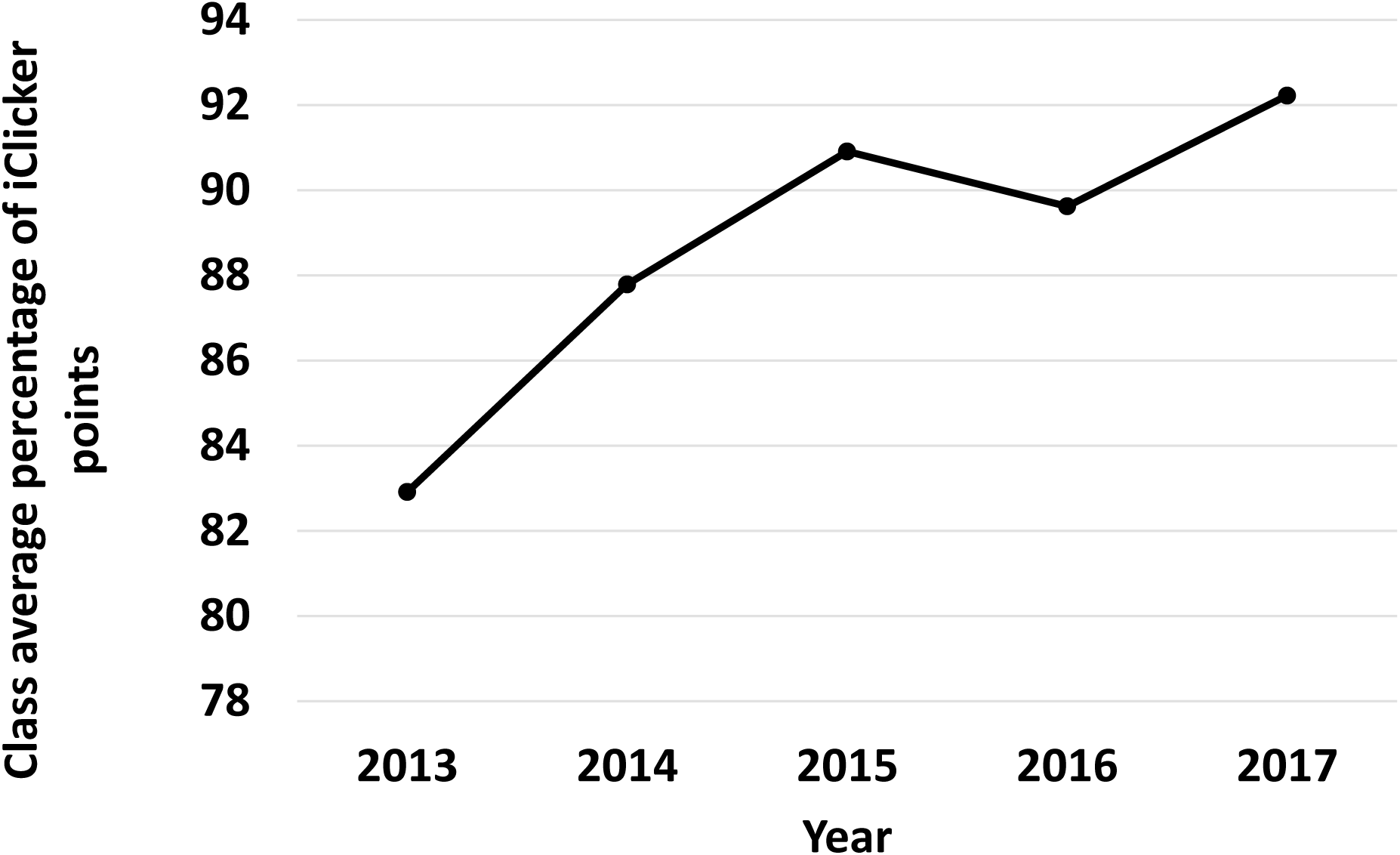
Student attendance. Student attendance was assessed by calculating the class average of iClicker points obtained during the entire semester. Points were assigned for every in-class question for which a response was registered irrespective of whether the response was correct or not.

**Fig. 5.**
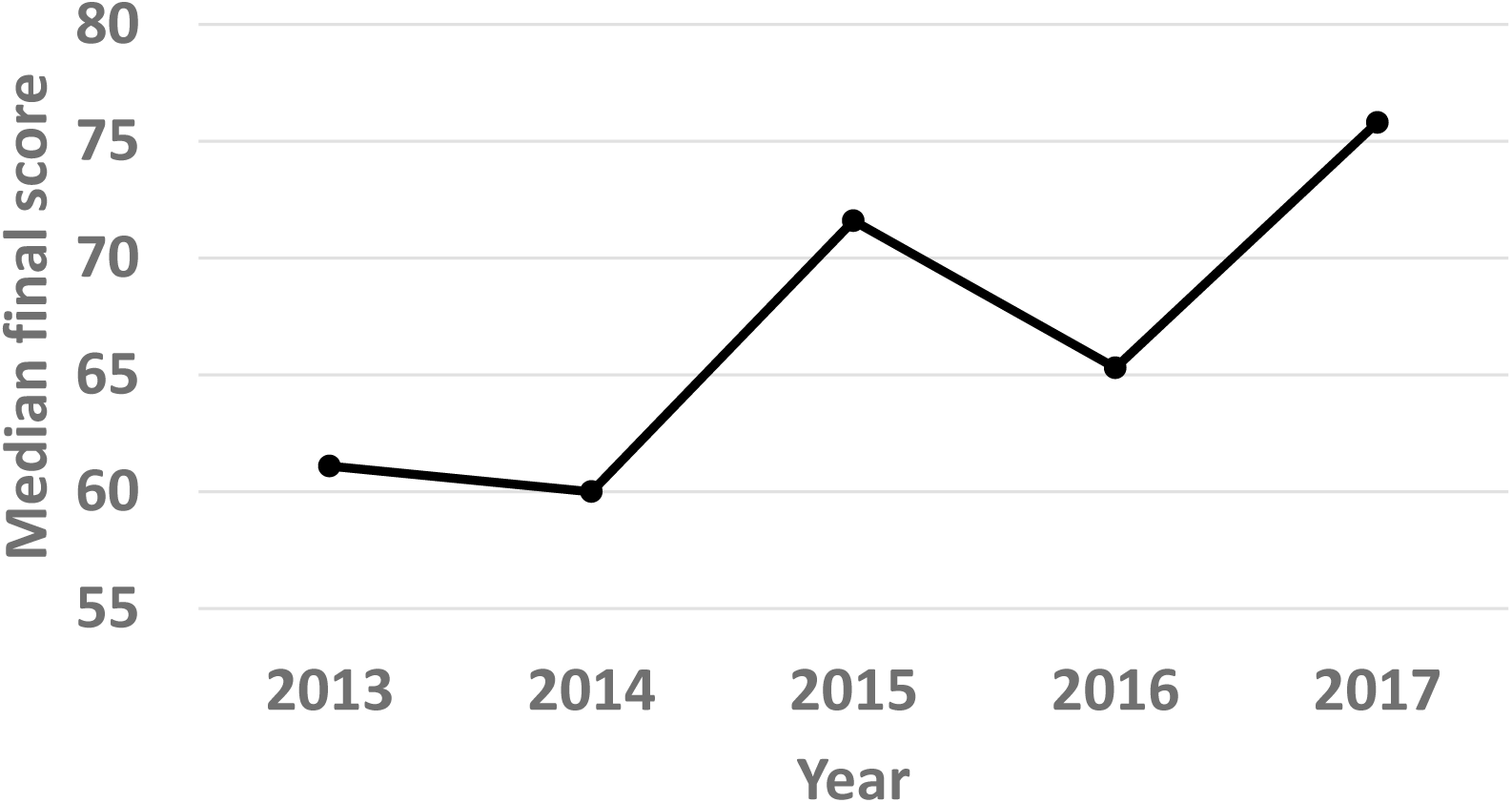
Comprehensive final analysis. The mean final score each year for the 95 question comprehensive final exam that covers each topic.

## Discussion

My analysis of course metrics from the years 2013-2017 indicates that in my large microbiology lecture course, lecturing with the chalk-talk method was far superior to a traditional PowerPoint lecture. The students preferred the chalk-talk approach, better retained key concepts, and they ultimately achieved higher scores. These five years are not a tightly controlled study as small differences were implemented each year to try and improve the course. For example, I suspect the increase in students receiving a 4.0 in 2017 was in part due to the audio of the lectures being made available to the class. But the general trend is clear in all data examined, and there is clearly a demarcation between the years when PowerPoint lecture was used versus the chalk-talk format.

I estimate that the course material was reduced around 40% when chalk-talk was used so the students in 2013 to 2014 were responsible for more material. This reduction was a conscious decision on my part as it had become clear that too much material was being covered. This reduction further highlights one of the main advantages of the chalk-talk approach. Using this approach forced me to focus on only the most important concepts and remove extraneous information. The reduction of material was not a removal of any of the core concepts, but rather a reduction in specific examples of a given topic. For example, rather than providing the students with four examples of sRNA regulation, I now only give them two. But understanding what sRNAs are and how they function remains a core concept of the course. This is evidenced by the fact that my lecture syllabus remained unchanged from 2013-2017. In essence, switching formats forced me to reduce the content of the course to the key microbial genetics concepts that I felt the students should learn, which is process that is being undertaken in all science fields as we continually expand our information base (15, 16). Moreover, the cumulative final exam scores increased when chalk-talk was used, providing evidence that key concepts were better retained.

Furthermore, an analysis of the SIRS scores does not suggest that the improvements to the course are entirely due to reduced course material. Importantly, the amount of information taught was identical in 2015-2017. If the improvements observed were simply due to a reduction in the amount of material, one would predict that SIRS results in 2015 would be similar in 2016 and 2017. However, this is not the case as improvements were observed in 2016 and 2017 compared to 2015, even though the amount of material did not change. Such a trend is also evident when examining specific SIRS comments (Fig. 3) and the final grades (Fig. 4) I speculate that these further improvements were due to my own increased experience with the chalk-talk format. This five-year study also highlights the important point that it is problematic to truly measure effectiveness of PowerPoint compared to chalk-talk by simply presenting one or a subset of lectures in each format and comparing the results. In order to gain the most benefit of the chalk-talk style, the course must be designed around this format and the instructor must be experienced using it as the largest benefits of using chalk-talk only became apparent only in 2017 (year 3). A strength of the data presented here, although it is not completely controlled, is that it spans multiple years allowing for a more real-world conclusion that accounts for instructor experience and course variation. Even though small changes were made to my course from year to year, the overall benefits that were observed upon switching to chalk-talk are quite clear. Whether a similar reduction of material when using PowerPoint would lead to similar gains is a question that requires further study.

There are several other factors to consider when comparing PowerPoint to chalk-talk. One important consideration of switching to chalk-talk is that research has shown seeing the lips and face during speech can promote increased comprehension, and if the instructor’s back is turned as during writing on a blackboard this cannot occur (17). However, by writing on a lap-top that was projecting on screens while facing the students overcomes this limitation. A large body of evidence indicates that actively taking notes, and later revisiting those notes for exam preparation contributes to increased learning (18-21). The chalk-talk approach I used certainly increased the amount of note-taking compared to using PowerPoint where the slides were provided before the lecture. On the other hand, excessive note-taking could lead to excessive demands on student’s cognitive function preventing learning during the lecture (19) and providing the slides beforehand could allow the students to spend less time note-taking leading to increased learning during class (22, 23). Although Babb and Ross found that providing lecture notes before class compared to after class increased participation and class attendance, there was no significant impact on exam scores (24). Whether or not chalk-talk approaches with partial or complete lecture notes outlines would improve performance over providing no lecture notes would be an interesting question to study further.

In the digital information age, it is challenging to maintain student engagement in large lecture courses with electronic devices offering a constant distraction. Banning these devices or disabling internet access in classrooms has been attempted as one solution to this problem. The data shown here clearly indicate that another solution is to use a chalk-talk approach, with limited PowerPoint slides for difficult diagrams, that requires active listening and note-taking without written lectures posted. My findings are consistent with other studies that conclude mixed lecture formats are the most effective lecturing styles (25-27). The data presented here show that when used appropriately, chalk-talk lecturing styles can be substantially more enjoyable and effective as the primary lecturing format in a large microbiology lecture course.

## Acknowledgements

I wish to thank Patrick Venta and John Merrill for helpful comments on the manuscript, and Macy Pell, Geoff Severin, Nico Fernandez, and Brian Hseuh for evaluating SIRS comments. I am grateful for the NSF-CAREER award MCB1253684, which encourages efforts at improving teaching and outreach, and the many wonderful Michigan State University microbial genetics students that are the basis for this research.

